# Crystal structure of Nsp15 endoribonuclease NendoU from SARS-CoV-2

**DOI:** 10.1101/2020.03.02.968388

**Authors:** Youngchang Kim, Robert Jedrzejczak, Natalia I. Maltseva, Michael Endres, Adam Godzik, Karolina Michalska, Andrzej Joachimiak

## Abstract

Severe Acute Respiratory Syndrome Coronavirus 2 is rapidly spreading around the world. There is no existing vaccine or proven drug to prevent infections and stop virus proliferation. Although this virus is similar to human and animal SARS- and MERS-CoVs the detailed information about SARS-CoV-2 proteins structures and functions is urgently needed to rapidly develop effective vaccines, antibodies and antivirals. We applied high-throughput protein production and structure determination pipeline at the Center for Structural Genomics of Infectious Diseases to produce SARS-CoV-2 proteins and structures. Here we report the high-resolution crystal structure of endoribonuclease Nsp15/NendoU from SARS-CoV-2 – a virus causing current world-wide epidemics. We compare this structure with previously reported models of Nsp15 from SARS and MERS coronaviruses.

## Introduction

Severe Acute Respiratory Syndrome Coronavirus 2 (SARS-CoV-2) is an etiologic agent responsible for the current outbreak of Coronavirus Disease 2019 (COVID-19). Over the past two months, the pathogen has infected over 88,000 people and caused at least 3,000 deaths. As we were preparing this report the numbers have doubled. Although currently mainly concentrated in China, the virus is spreading worldwide rapidly and is found in 64 countries and all continents (www.trackcorona.live). Millions of people are being quarantined and the epidemics impacts the world economy. There is no existing vaccine or proven drug for this disease, but various treatment options, for example utilizing medicines effective in other viral ailments, are being attempted.

Coronaviruses are enveloped, non-segmented positive-sense RNA viruses from the order nidoviruses that have the largest genome among RNA viruses. The genome contains a large replicase gene encompassing nonstructural proteins (Nsps), followed by structural and accessory genes. Due to ribosomal frameshifting, the replicase gene encodes two ORFs, rep1a and rep1b, that are translated into two large polyproteins, pp1a and pp1ab (Cui *et al.*, 2019). These polypeptides are processed by two viral proteases: 3C-like protease (3CLpro, encoded by Nsp5), and papain-like protease (PLP, encoded within Nsp3). The cleavage yields 16 viral Nsps (Baez-Santos *et al.*, 2015) that assemble into a large membrane-bound replicase complex and exhibits multiple enzymatic activities. While several functions of Nsps have been linked to RNA replication and processing of subgenomic RNAs, the roles of some proteins are poorly understood or remain unknown.

One of such enigmatic enzymes, corresponding to Nsp15, is a nidoviral RNA uridylate-specific endoribonuclease (NendoU) carrying C-terminal catalytic domain belonging to the EndoU family. EndoU enzymes are present in all kingdoms of life, where they play various biological functions associated with RNA processing. All characterized family members display an RNA endonuclease activity producing 2’-3’ cyclic phosphodiester and 5’-hydroxyl termini (Ulferts & Ziebuhr, 2011). The viral and eukaryotic enzymes act on both, single- and double stranded RNA and are specific for uridine. The prototypic member of the family was discovered in *Xenopus laevis*, and thus named XendoU, where it is associated with maturation of intron-encoded small nucleolar RNAs (snoRNA) (Caffarelli *et al.*, 1994, Caffarelli *et al.*, 1997, Gioia *et al.*, 2005, Laneve *et al.*, 2003). The human homolog, placental protein 11 (PP11, HendoU) plays yet an unknown role in the placental tissue (Laneve *et al.*, 2008), but is also expressed in cancer cells. In *Drosophila*, DendoU has been found to be relevant for nervous system physiology and pathology (Laneve *et al.*, 2017). The EndoU representatives are also present in prokaryotes, with the best characterized enzyme lacking uridylate specificity of the family and acting as a tRNAse toxin in bacterial communication (Michalska *et al.*, 2018).

In viruses, the NendoU protein is conserved among coronaviruses, arteriviruses and toroviruses, but is absent in non-vertebrate - infecting representatives of the nidoviruses order: mesoniviruses and roniviruses. While initially Nsp15 was thought to directly participate in viral replication, it was later shown that Nsp15-deficient coronaviruses were viable and replicating, rising doubts about the enzyme role in that process. More recently, it was proposed that NendoU activity of Nsp15 is responsible for the protein interference with the innate immune response (Deng *et al.*, 2017) though other studies indicate that the process is independent of the endonuclease activity (Liu *et al.*, 2019). Nevertheless, Nsp15 is essential in coronavirus biology.

Nsp15s from coronaviruses share little sequence similarity with their arteriviruses counterpart Nsp11: the SARS-CoV Nsp15 has only 22% identical residues with Porcine Reproductive and Respiratory Syndrome virus (PRRSV) Nsp11. Within arteriviruses alone sequence conservation of Nsp11 is generally low, while Nsp15 of coronaviruses are more conserved, with the current SARS-CoV-2 sharing 88% sequence identity and 95% similarity with its known closest homolog from SARS-CoV. Nsp15 from Middle East Respiratory Syndrome coronavirus (MERS-CoV) is more distant, with ~50% sequence identity and ~65% similarity and H-CoV-229E being even more distant with 43% sequence identity and 56% similarity.

After the 2002-2003 SARS outbreak there was an increased interest in virus-encoded endoribonucleases. The first two structures of Nsp15 were determined from Mouse Hepatitis Virus (MHV) (PDB id 2GTH, 2GTI (Xu *et al.*, 2006)) and SARS coronavirus (PDB id 2H85, (Ricagno *et al.*, 2006). Currently, there are in total eight structures available for Nsp15 including additional two from SARS-CoV (PDB ids: 2OZK, 2RHB, (Joseph *et al.*, 2007)), one from MERS-CoV (PDB id 5YVD, (Zhang *et al.*, 2018)) and two from human coronavirus H-CoV-229E (PDB ids 4S1T, 4RS4, (Huo & Liu, 2015)). Among Nsp11 endoribonucleases, there are two structures determined of PRRSV (Porcine Reproductive and Respiratory Syndrome Virus) (PDB ids 5EYI, 5DA1, (Zhang *et al.*, 2017)), and two for EAV (Equine Arteritis Virus, PDB ids 5HC1, 5HBZ, (Zhang *et al.*, 2017)).

The structural studies of SARS- and MERS-CoV Nsp15s showed that the protein forms hexamers made of dimers of trimers. The 39 kDa monomeric unit, composed of ~345 residues, folds into three domains: N-terminal, middle domain and C-terminal catalytic NendoU domain.

Rapid upsurge and proliferation of SARS-CoV-2 raised questions about how this virus could became so much more transmissible as compared to the SARS and MERS coronaviruses. The proteins of the coronavirus are mutating, although not very fast, but these changes may contribute to the virus virulence. Here, we report the first crystal structure of SARS-CoV-2 Nsp15 at 2.20 Å resolution. The Nsp15 structure is very similar to the SARS-CoV and MERS-CoV but it shows some differences that may contribute to SARS-CoV-2 virulence.

## Results and Discussion

### Protein production and structure determination

We have used synthetic gene of SARS-CoV-2 coronavirus *nsp15*, which was codon optimized for expression in *E. coli*, to produce soluble protein. Nsp15 was purified and crystallized exercising the well-established structural genomics pipeline. The structure was determined by molecular replacement and refined to 2.20 Å resolution, as described in Materials and Methods and Table 1. The entire process from protein expression to structure deposition to PDB took 5 days.

**Table 1.**
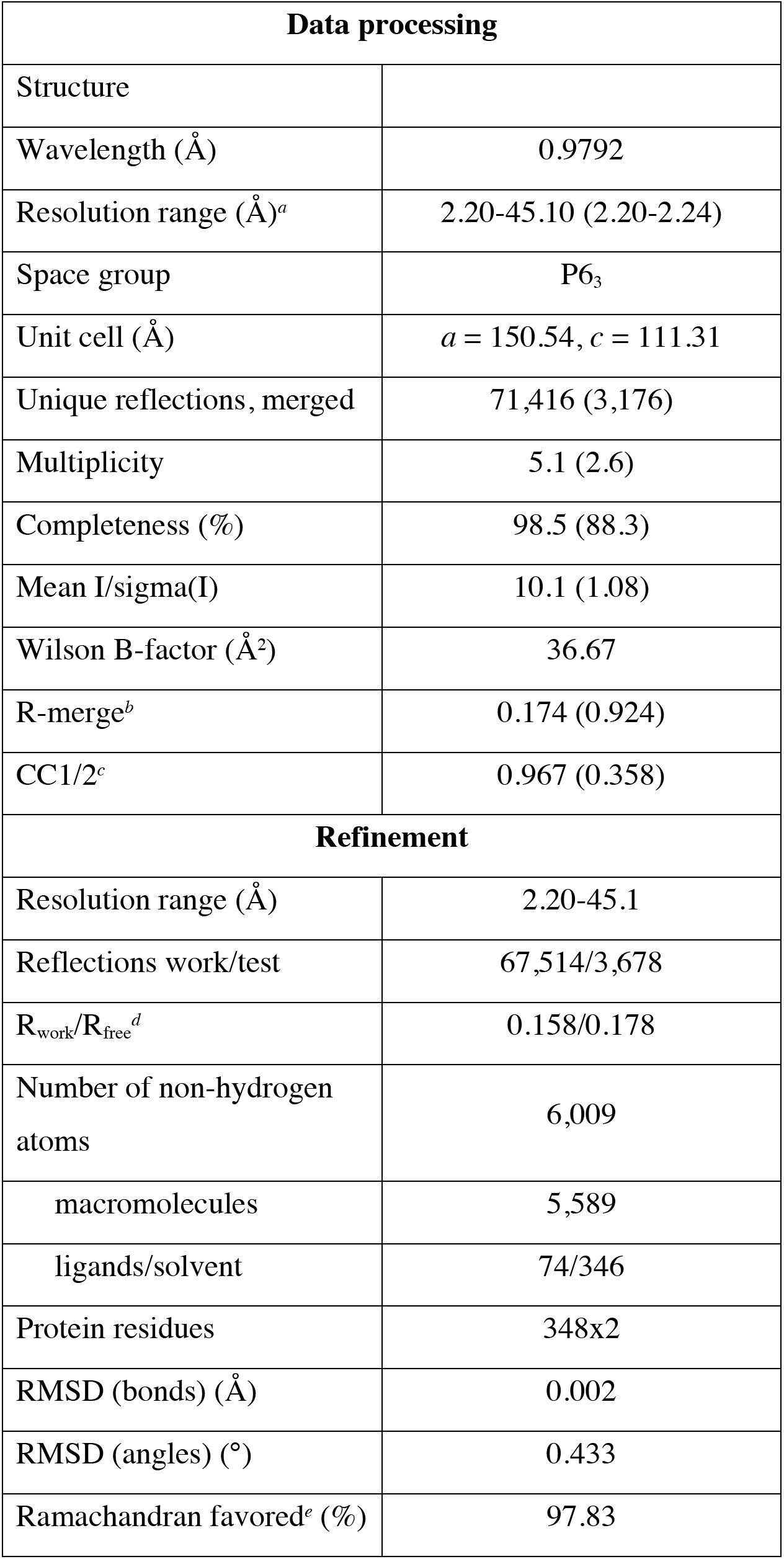
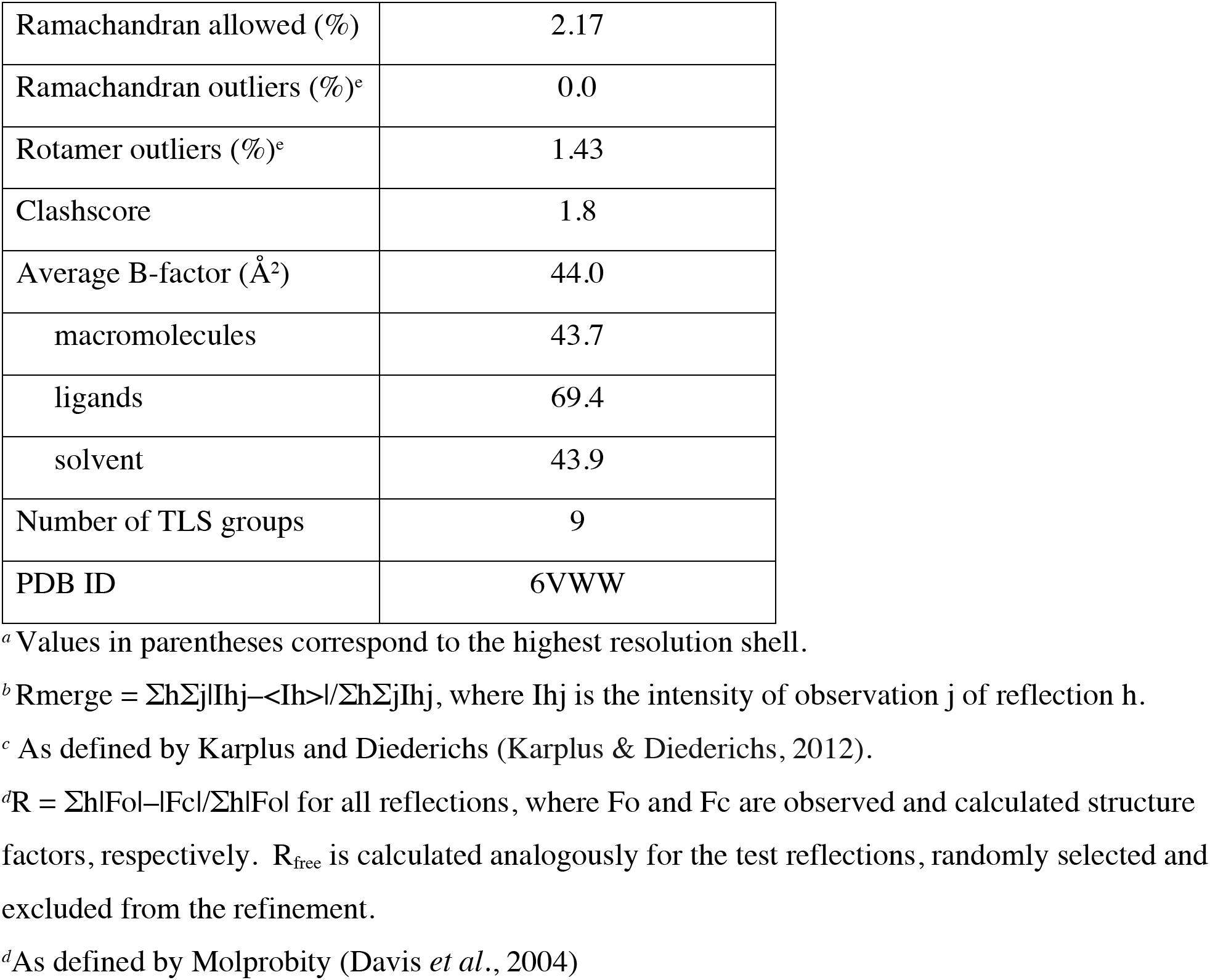
Data processing and refinement statistics.

The structure of SARS-CoV-2 coronavirus Nsp15 is of good quality and it refined to crystallographic R_work_ of 15.8% and R_free_ of 17.8%. The asymmetric unit contains two monomers of the Nsp15 protein. The electron density map is of high quality throughout the structure and the model covers the sequence from M1 to Q347. Currently, this is the most complete and the highest resolution structure of coronavirus Nsp15. 346 water molecules, eight glycerol molecules, three acetate ions, one magnesium and one chloride ion were identified in the electron density maps.

### Overall structure

The structure of SARS-CoV-2 Nsp15 monomer is very similar to other Nsp15s from coronaviruses (Fig. 1 and 2). It features three distinct domains. The N-terminal domain is composed of an antiparallel β-sheet (strands β1, β2, and β3) wrapped around two α-helices (α1 and α2). The subsequent middle domain is formed by 10 β-strands organized in three β-hairpins (β5-β6, β7-β8, β12-β13), a mixed β-sheet (β4, β9, β10, β11, β14, β15) and three short helices, two α and one 3_10_ (α3, η4, and α5). The C-terminal catalytic NendoU domain contains two antiparallel β-sheets (β16-β17-β18, β19-β20-β21) with their edges hosting a catalytic site. β18, even though is broken in the secondary structure annotation, is treated here as a single element. The concave surface of the β-sheets is flanked by five α-helices (α6, α7, α8, α9, α10).

**Figure 1.**
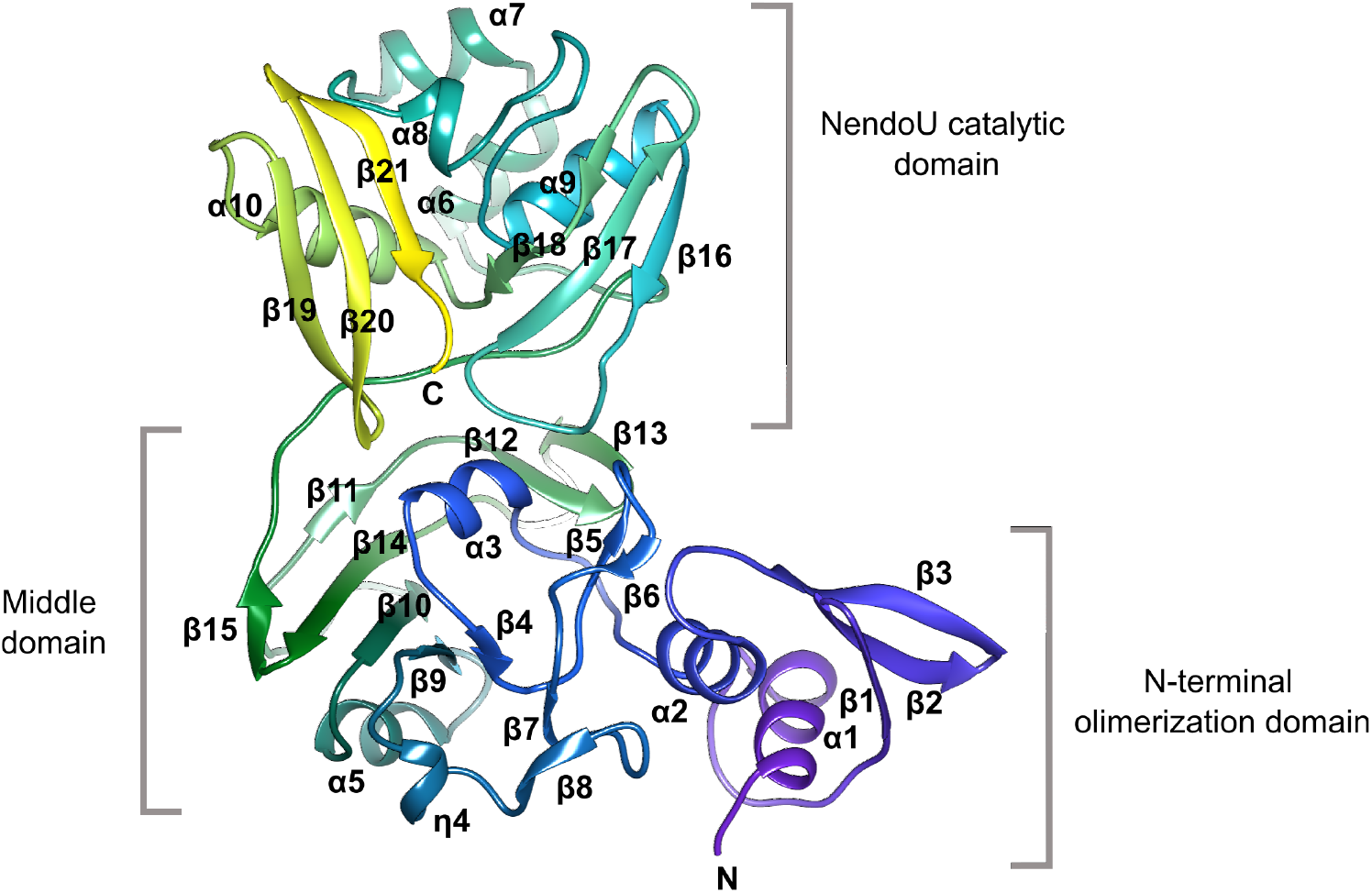
The structure of SARS-CoV-2 monomer.

**Figure 2.**
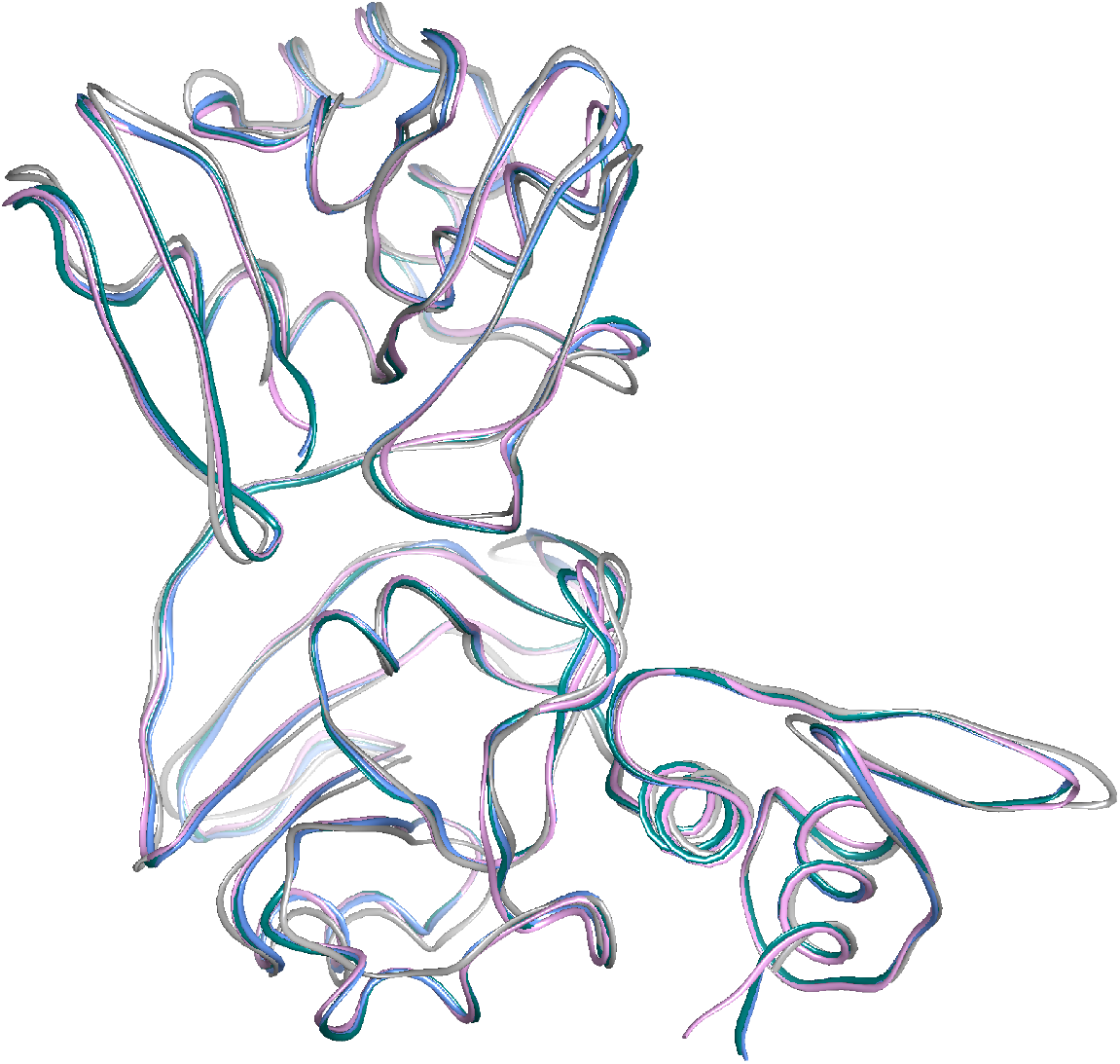
Superposition of Nsp15 monomers: SARS-CoV-2 (teal, chain A; blue, chain B), SARS-CoV (pink), MERS-CoV (grey).

As expected from high sequence identity (Fig. 3), the structure aligns best with SARS-CoV Nsp15 (0.47 Å RMSD of chain A with PDB id: 2H85, chain A; sequence identity 88%), and also shows good agreement with Nsp15 from MERS-CoV (1.17 Å RMSD of chain A with PDB id 5YVD, chain A; sequence identity 51%). The structural homology is not only observed in positions of α-helices and β-strands but surprisingly in several loop regions (for example the conformation of loop L1 (F16 – P24) is virtually identical in all three proteins). But there are some interesting differences between SARS- and MERS-CoVs proteins in loop regions (β8-β9, β10-β11) with the largest (more than 5 Å) centering on the loop between strand β13 and β14 where MERS-CoV Nsp15 has three-residues insertion. These relatively small changes in the subunit structures translate into larger shifts in the hexamer (see below).

**Figure 3.**
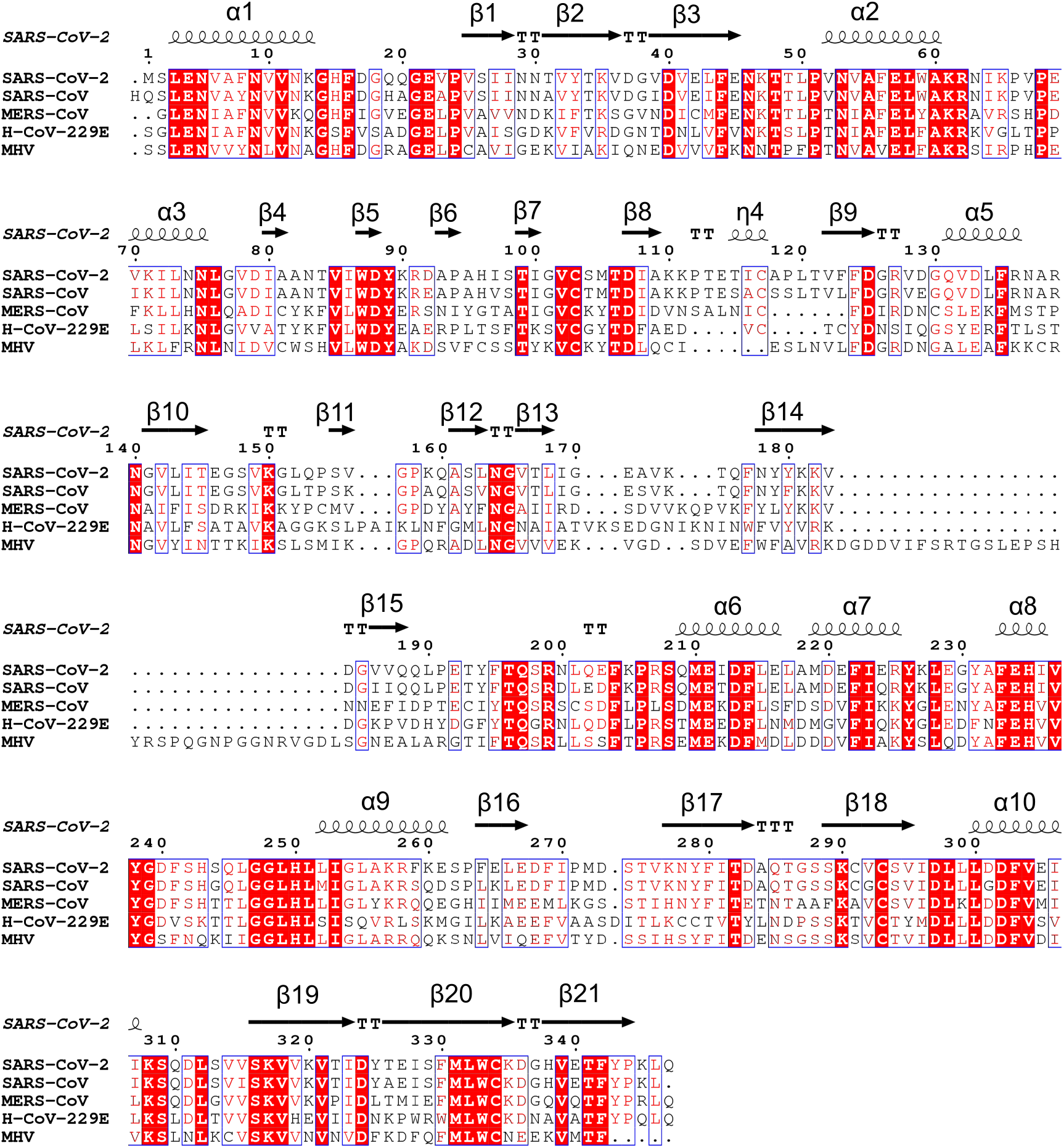
Sequence alignment of SARS-CoV-2 Nsp15 coronaviral homologs with structures available in the PDB: SARS-CoV-2 (6VWW), SARS-CoV (2H85), MERS-CoV (5YVD), H-CoV-29E (4RS4) MHV (2GTH). The secondary structure elements labeled for SARS-CoV-2 Nsp15.

The SARS-CoV-2 NendoU monomers assemble into a double-ring hexamer, generated by a dimer of trimers (Fig. 4). This is in agreement with PISA calculations, estimating trimers to be a stable form of the enzyme, and the previous experimental work capturing SARS-CoV Nsp15 trimeric intermediates (Guarino *et al.*, 2005). It has been shown that hexamer is essential for the enzymatic activity. A ~100 Å long and narrow (10-15 Å diameter) channel runs down the 3-fold axis. This channel is accessible to solvent from the top, bottom and also through three side openings in the middle of hexamer. The hexamer is stabilized by the interactions of N-terminal oligomerization domains, but also each subunit domain contributes to oligomer interface. As a result, monomers interact extensively with all five other subunits of the hexamer, making the hexamer potentially very sensitive to mutations that can disrupt oligomeric assembly (Fig. 4). The middle domains are the most transposed out in the hexamer creating the concave surfaces that may serve as an interaction hubs.

**Figure 4.**
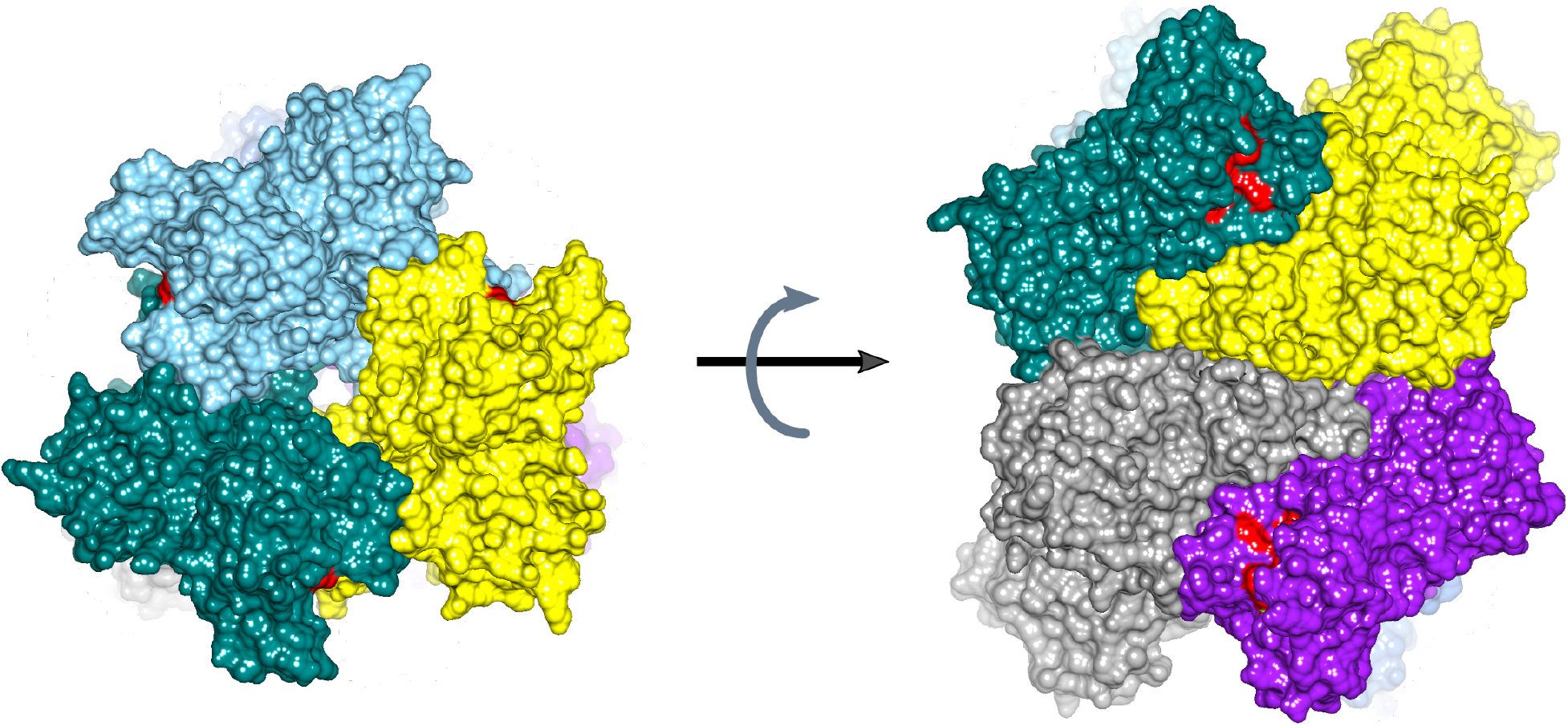
**Structure of SARS-CoV-2 hexamer** in surface representation with each subunit shown in different color. The active site residues are colored red.

The SARS-CoV-2 NendoU oligomer resembles those of SARS-CoV, H-CoV-229E and MERS-CoV enzymes. As with the monomeric folds, the SARS-CoV-2 hexamer shows higher similarity to the SARS-CoV assembly than to H-CoV-229E and MERS-CoV. The largest difference between SARS-CoV-2 and SARS-CoV seems to occur in the position of middle domains. The differences with H-CoV-229E are still more significant and show shifts in positions of α-helices, β-sheets and loops. Similarly, MERS-CoV enzyme show significant structural changes, particularly in the loops of the middle domain. This suggests that the SARS-CoV-2 enzyme operates most likely a manner very similar to SARS-CoV, H-CoV-229E and MERS CoV homologs, though it still may display different catalytic properties and potentially altered substrate specificity.

### NendoU active site

Recombinant Nsp15s were shown to have Mn^+2^-dependent endoribonuclease activity that cut double-stranded (ds) RNA substrates with specificity towards uridylate in unpaired regions (Bhardwaj *et al.*, 2004, Bhardwaj *et al.*, 2008, Bhardwaj *et al.*, 2006, Ivanov *et al.*, 2004). While the early work suggested cleavage upstream and downstream of U (Bhardwaj *et al.*, 2004, Ivanov *et al.*, 2004), the 2006 more detailed study demonstrated the reaction occurs on the 3’ end of U (Bhardwaj *et al.*, 2006). The enzyme carries out transesterification reaction releasing 2’-3’cyclic phosphate end (Ivanov *et al.*, 2004), as has been demonstrated for other members of EndoU family (Nedialkova *et al.*, 2009, Renzi *et al.*, 2006), though some studies report subsequent hydrolysis of the cyclic compound (Nedialkova *et al.*, 2009).

The catalytic function of Nsp15 resides in the C-terminal NendoU domain. The active site, located in a shallow groove between the two β-sheets, carries six key residues conserved among SARS-CoV-2, SARS-CoV and MERS-CoV proteins: His235, His250, Lys290, Thr341, Tyr343, and Ser294 (Fig. 5). The main chain architecture of this region as well as side chain conformations of the active site residues (with exception of Lys290) are conserved between all three proteins. The two histidine residues are contributed by the helical layer of the domain, while pairs lysine/serine and threonine/tyrosine originate from two β-strands representing edges of the β-sheets. His235, His250, Lys290 have been proposed to constitute the catalytic triad, based on the similarity of their mutual arrangement to the active site of ribonuclease A (Ricagno *et al.*, 2006). In this scenario, His235 plays a role of the general acid while His250 acts a base. By the same reasoning, Ser294 together with Tyr343 are believed to govern U specificity, resembling the roles of Phe120 and Thr45 (the B1 subsite specific for pyrimidine base) in RNAse A in base recognition (delCardayre & Raines, 1994). Specifically, Ser294 analogs have been proposed to interact with the carbonyl oxygen atom O2 of uracil via the main chain nitrogen atom, while the hydroxyl group would bind to the nitrogen atom ((Bhardwaj *et al.*, 2004, Bhardwaj *et al.*, 2008, Bhardwaj *et al.*, 2006, Ivanov *et al.*, 2004). Notably though, not all mutational data support the hypothesis of the discriminatory role of Ser and, as an alternative, Thr341 has been also considered (Ricagno *et al.*, 2006).

**Figure 5.**
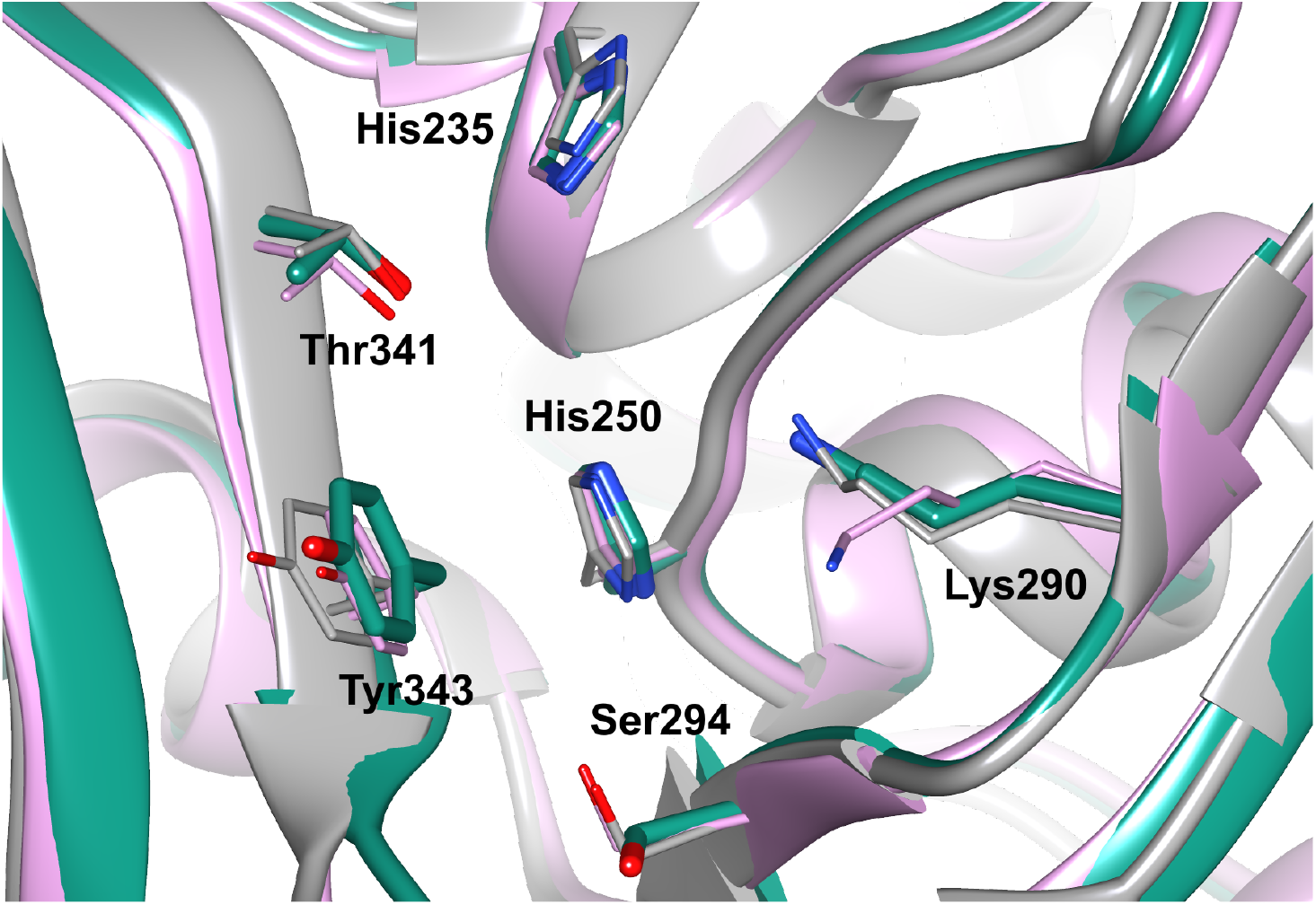
Superposition of the SARS-CoV-2 Nsp15 NendoU active site with its homologs: SARS-CoV-2 (teal, chain A), SARS-CoV (pink), MERS-CoV (grey).

The Mn^+2^ dependence has been observed for most EndoU members, but for example Nsp11 does not show such behavior (Nedialkova *et al.*, 2009), and appears to be a common feature of NendoU subfamily. However, the metal binding site was never located, though it is important to note that there is no structure of the protein/RNA complex. In our structure, there are two subunits in the asymmetric unit. One of them has an electron density peak near the active site that may correspond to metal ion and we tentatively modeled it as a magnesium ion, although the coordination sphere is poor. The metal ion is coordinated by oxygen atom of carboxylate Asp283 (2.61 Å), hydroxyl group of Ser262 (2.08 Å) and main chain carbonyl oxygen atom of Pro263 (2.28 Å). Nearby, there is a side chain of Arg258. If this would be a manganese ion, this arginine could coordinate the metal ion using nitrogen of guanidinium moiety (distance to magnesium ion is 4.1 Å). These residues are conserved in SARS but not in MERS enzymes. We propose that this is the metal binding site required for maintaining conformation of the active site and substrate during catalysis.

In the hexameric context, the active sites are located on the top and bottom of the assembly. NendoU binds single and double stranded RNA and large substrate can access these six sites from a site of the hexamer. All six sites can be occupied simultaneously. Due to extensive interactions between protein subunits, there is possibility for cooperativity or anti-cooperativity between binding sites. By analogy to SARS-CoV Nsp15 and SARS-CoV-2 Nsp15, the hexamer most likely represents an active form of the enzyme.

## Conclusions

We have determined the high-resolution crystal structure of endoribonuclease NendoU from SARS-CoV-2. The structure is homologous to SARS- and MERS-CoVs Nsp15s and shows a hexamer, the functionally active form of the endoribonuclease. The active site residues are conserved both in terms of sequence and conformation. The structural comparisons suggest that inhibitors of SARS-CoV Nsp15 have good chance to inhibit also the SARS-CoV-2 homolog but inhibitors of MERS-CoV NendoU are unlikely to inhibit the enzyme.

## Materials and methods

### Gene cloning, protein expression and purification

The gene cloning, protein expression and purification were performed as reported previously (Makowska-Grzyska *et al.*, 2014). Briefly, the gene for Nsp15 SARS-CoV-2 was optimized for *E. coli* expression using the OptimumGene codon optimization algorithm followed by manual editing and then synthesized cloned directly into pMCSG53 vector (Twist Bioscience). The plasmid was transformed into the *E. coli* BL21(DE3)-Gold strain (Stratagene). For large-scale purification of the protein, a 4 L culture of LB Lennox medium was grown at 37°C (190 rpm) in presence of ampicillin 150 μg/ml. Once the culture reached OD_600_ ~1.0, the temperature setting was changed to 4°C. When bacterial suspension cooled down to 18°C it was supplemented with the following components to indicated concentration: 0.2 mM IPTG, 0.1% glucose, 40mM K_2_HPO_4_. The temperature was set to 18°C for 20 hours incubation. Bacterial cells were harvested by centrifugation at 7,000*g* and cell pellets were resuspended in a 12.5 ml lysis buffer (500 mM NaCl, 5% (v/v) glycerol, 50 mM HEPES pH 8.0, 20 mM imidazole and 10 mM β-mercaptoethanol) per liter culture and sonicated at 120W for 5 minutes (4 sec ON, 20 sec OFF). The cellular debris was removed by centrifugation at 30,000*g* for one hour at 4°C. Supernatant was mixed with 4 ml of Ni^2+^ Sepharose (GE Healthcare Life Sciences) equilibrated with lysis buffer supplemented to 50 mM imidazole pH 8.0 and suspension was applied on Flex-Column (420400-2510) connected to Vac-Man vacuum manifold (Promega). Unbound proteins were washed out via controlled suction with 160 ml of lysis buffer (50 mM imidazole). Bound proteins were eluted with 20 ml of lysis buffer supplemented to 500 mM imidazole pH 8.0. 2 mM DTT was added followed by Tobacco Etch Virus (TEV) protease treatment at 1:20 protease:protein ratio. The solution was left at 4°C overnight. Unfortunately, for this particular construct, TEV protease was not able to cleave off the His tag. Nsp15 was successfully separated from TEV protease on Superdex 200 column equilibrated in lysis buffer where 10 mM β-mercaptoethanol was replaced by 1 mM TCEP. Fractions containing Nsp15 were collected. Lysis buffer was replaced on 30 kDa MWCO filter (Amicon-Millipore) via 10X concentration/dilution repeated 3 times to crystallization buffer (150 mM NaCl, 20 mM HEPES pH 7.5, 1 mM TCEP). Final concentration of Nsp15 was 36 mg/ml.

### Crystallization

Crystallization experiments also were conducted as described previously, with slight modifications. The sitting-drop vapor-diffusion method was used with the help of the Mosquito liquid dispenser (TTP LabTech) in 96-well CrystalQuick plates (Greiner Bio-One). Crystallizations were performed with the protein-to-matrix ratio of 1:1. MCSG4 (Anatrace), SaltRX (Hampton) and INDEX (Hampton) screens were used for protein crystallization at 16°C. The best conditions were MCSG4 H11 (0.2 M calcium acetate, 0.1 M HEPES/NaOH pH 7.5, 10% PEG 8000). Diffraction-quality crystals of Nsp15 suitable for data collection appeared after 12 hours.

### Data collection, structure determination and refinement

Prior to data collection at 100 K, all cryoprotected crystals of Nsp15 were flash-cooled in liquid nitrogen. The x-ray diffraction experiments were carried out at the Structural Biology Center 19-ID beamline at the Advanced Photon Source, Argonne National Laboratory. The diffraction images were recorded on the PILATUS3 6M detector K using 0.5° rotation and 0.5 sec exposure for 100°. The data set was processed and scaled with the HKL3000 suite (Minor *et al.*, 2006). Intensities were converted to structure factor amplitudes in the Ctruncate program (French & Wilson, 1978, Padilla & Yeates, 2003) from the CCP4 package (Winn *et al.*, 2011). The structure was determined using molrep (Vagin & Teplyakov, 2010) implemented in the HKL3000 software package and SARS-CoV Nsp15 structure (PDB id 2H85) as a search model. The initial solution was manually adjusted using COOT (Emsley & Cowtan, 2004) and then iteratively refined using COOT, PHENIX (Adams *et al.*, 2010) and REFMAC (Murshudov *et al.*, 1997, Winn *et al.*, 2011). Initially REFMAC refinement worked better for the structure which was subsequently moved to PHENIX to finalize the refinement. Throughout the refinement, the same 5% of reflections were kept out throughout from the refinement (in both REFMAC and PHENIX refinement). The final structure converged to a good R_work_ = 0.158 and R_free_ = 0.178 with regards to data quality. The stereochemistry of the structure was checked with PROCHECK (Laskowski *et al.*, 1993) and the Ramachandran plot and validated with the PDB validation server. The data collection and processing statistics are given in Table 1. The atomic coordinates and structure factors have been deposited in the Protein Data Bank under accession code 6VWW. As we were preparing this report we have determined and deposited to PDB higher resolution 1.90 Å structure of the SARS-CoV-2 Nsp15 in complex with citrate (PDB id 6W01). This structure will be released to public domain immediately after processing and will be discussed in a future publication.

## Acknowledgements

We truthfully thank the members of the SBC at Argonne National Laboratory, especially Darren Sherrell and Alex Lavens for their help with setting beamline and data collection at beamline 19-ID. We thank Lukasz Jaroszewski and Monica Rosas Lemus for construct design. Funding for this project was provided in part by federal funds from the National Institute of Allergy and Infectious Diseases, National Institutes of Health, Department of Health and Human Services, under Contract HHSN272201700060C. The use of SBC beamlines at the Advanced Photon Source is supported by the U.S. Department of Energy (DOE) Office of Science and operated for the DOE Office of Science by Argonne National Laboratory under Contract No. DE-AC02-06CH11357.

## References

Adams, P. D., Afonine, P. V., Bunkoczi, G., Chen, V. B., Davis, I. W., Echols, N., Headd, J. J., Hung, L. W., Kapral, G. J., Grosse-Kunstleve, R. W., McCoy, A. J., Moriarty, N. W., Oeffner, R., Read, R. J., Richardson, D. C., Richardson, J. S., Terwilliger, T. C. & Zwart, P. H. (2010). Acta Crystallogr., Sect D: Biol. Crystallogr. 66, 213–221.

Baez-Santos, Y. M., St John, S. E. & Mesecar, A. D. (2015). Antiviral Res. 115, 21–38.

Bhardwaj, K., Guarino, L. & Kao, C. C. (2004). J Virol 78, 12218–12224.

Bhardwaj, K., Palaninathan, S., Alcantara, J. M., Yi, L. L., Guarino, L., Sacchettini, J. C. & Kao, C. C. (2008). J Biol Chem 283, 3655–3664.

Bhardwaj, K., Sun, J., Holzenburg, A., Guarino, L. A. & Kao, C. C. (2006). J Mol Biol 361, 243–256.

Caffarelli, E., Arese, M., Santoro, B., Fragapane, P. & Bozzoni, I. (1994). Mol Cell Biol 14, 2966–2974.

Caffarelli, E., Maggi, L., Fatica, A., Jiricny, J. & Bozzoni, I. (1997). Biochem Biophys Res Commun 233, 514–517.

Cui, J., Li, F. & Shi, Z. L. (2019). Nat. Rev. Microbiol. 17, 181–192.

Davis, I. W., Murray, L. W., Richardson, J. S. & Richardson, D. C. (2004). Nucleic Acids Res. 32, W615–619.

delCardayre, S. B. & Raines, R. T. (1994). Biochemistry 33, 6031–6037.

Deng, X., Hackbart, M., Mettelman, R. C., O’Brien, A., Mielech, A. M., Yi, G., Kao, C. C. & Baker, S. C. (2017). Proc Natl Acad Sci U S A 114, E4251–E4260.

Emsley, P. & Cowtan, K. (2004). Acta Crystallogr., Sect D: Biol. Crystallogr. 60, 2126–2132.

French, S. & Wilson, K. (1978). Acta Crystallogr., Sect. A: Found. Crystallogr. 34, 517–525.

Gioia, U., Laneve, P., Dlakic, M., Arceci, M., Bozzoni, I. & Caffarelli, E. (2005). J Biol Chem 280, 18996–19002.

Guarino, L. A., Bhardwaj, K., Dong, W., Sun, J., Holzenburg, A. & Kao, C. (2005). J Mol Biol 353, 1106–1117.

Huo, T. & Liu, X. (2015). Acta Crystallogr F Struct Biol Commun 71, 1156–1160.

Ivanov, K. A., Hertzig, T., Rozanov, M., Bayer, S., Thiel, V., Gorbalenya, A. E. & Ziebuhr, J. (2004). Proc Natl Acad Sci U S A 101, 12694–12699.

Joseph, J. S., Saikatendu, K. S., Subramanian, V., Neuman, B. W., Buchmeier, M. J., Stevens, R. C. & Kuhn, P. (2007). J Virol 81, 6700–6708.

Karplus, P. A. & Diederichs, K. (2012). Science 336, 1030–1033.

Laneve, P., Altieri, F., Fiori, M. E., Scaloni, A., Bozzoni, I. & Caffarelli, E. (2003). J Biol Chem 278, 13026–13032.

Laneve, P., Gioia, U., Ragno, R., Altieri, F., Di Franco, C., Santini, T., Arceci, M., Bozzoni, I. & Caffarelli, E. (2008). J. Biol. Chem. 283, 34712–34719.

Laneve, P., Piacentini, L., Casale, A. M., Capauto, D., Gioia, U., Cappucci, U., Di Carlo, V., Bozzoni, I., Di Micco, P., Morea, V., Di Franco, C. A. & Caffarelli, E. (2017). Sci Rep 7, 41559.

Laskowski, R. A., MacArthur, M. W., Moss, D. S. & Thornton, J. M. (1993). Journal of Applied Crystallography 26, 283–291.

Liu, X., Fang, P., Fang, L., Hong, Y., Zhu, X., Wang, D., Peng, G. & Xiao, S. (2019). Mol Immunol 114, 100–107.

Makowska-Grzyska, M., Kim, Y., Maltseva, N., Li, H., Zhou, M., Joachimiak, G., Babnigg, G. & Joachimiak, A. (2014). Methods Mol Biol 1140, 89–105.

Michalska, K., Quan Nhan, D., Willett, J. L. E., Stols, L. M., Eschenfeldt, W. H., Jones, A. M., Nguyen, J. Y., Koskiniemi, S., Low, D. A., Goulding, C. W., Joachimiak, A. & Hayes, C. S. (2018). Mol Microbiol 109, 509–527.

Minor, W., Cymborowski, M., Otwinowski, Z. & Chruszcz, M. (2006). Acta Crystallogr., Sect D: Biol. Crystallogr. 62, 859–866.

Murshudov, G. N., Vagin, A. A. & Dodson, E. J. (1997). Acta Crystallogr., Sect D: Biol. Crystallogr. 53, 240–255.

Nedialkova, D. D., Ulferts, R., van den Born, E., Lauber, C., Gorbalenya, A. E., Ziebuhr, J. & Snijder, E. J. (2009). J Virol 83, 5671–5682.

Padilla, J. E. & Yeates, T. O. (2003). Acta Crystallogr., Sect D: Biol. Crystallogr. 59, 1124–1130.

Renzi, F., Caffarelli, E., Laneve, P., Bozzoni, I., Brunori, M. & Vallone, B. (2006). Proc Natl Acad Sci U S A 103, 12365–12370.

Ricagno, S., Egloff, M. P., Ulferts, R., Coutard, B., Nurizzo, D., Campanacci, V., Cambillau, C., Ziebuhr, J. & Canard, B. (2006). Proc Natl Acad Sci U S A 103, 11892–11897.

Ulferts, R. & Ziebuhr, J. (2011). RNA Biol 8, 295–304.

Vagin, A. & Teplyakov, A. (2010). Acta Crystallogr., Sect D: Biol. Crystallogr. 66, 22–25.

Winn, M. D., Ballard, C. C., Cowtan, K. D., Dodson, E. J., Emsley, P., Evans, P. R., Keegan, R. M., Krissinel, E. B., Leslie, A. G., McCoy, A., McNicholas, S. J., Murshudov, G. N., Pannu, N. S., Potterton, E. A., Powell, H. R., Read, R. J., Vagin, A. & Wilson, K. S. (2011). Acta Crystallogr., Sect D: Biol. Crystallogr. 67, 235–242.

Xu, X., Zhai, Y., Sun, F., Lou, Z., Su, D., Xu, Y., Zhang, R., Joachimiak, A., Zhang, X. C., Bartlam, M. & Rao, Z. (2006). J Virol 80, 7909–7917.

Zhang, L., Li, L., Yan, L., Ming, Z., Jia, Z., Lou, Z. & Rao, Z. (2018). J Virol 92.

Zhang, M., Li, X., Deng, Z., Chen, Z., Liu, Y., Gao, Y., Wu, W. & Chen, Z. (2017). J Virol 91.

